# Neural Activity Dynamics in Primate Cortex Across Consciousness Levels: Insights from High-Density Neuropixel Recording

**DOI:** 10.1101/2025.10.09.681481

**Authors:** Majid Khalili-Ardali, Abhilash Dwarakanath, Maxime Roustan, Béchir Jarraya, Theofanis Panagiotaropoulos

## Abstract

This study investigates the anesthesia mechanisms induced by sevoflurane and how it modulates neural activity in the posterior parietal cortex (PPC) and prefrontal cortex (PFC) in Non Human Primates (NHPs) using high density Neuropixel probes. Spiking and local field potentials (LFPs) were recorded in two macaque monkeys under going four sevoflurane concentrations (2%, 3%, 4%, and 6%). We aimed to (i) quantify the emergence of anesthesia-induced Up/Down state dynamics, (ii) track changes in oscillatory power and inter-regional synchrony, and (iii) determine whether frontal and parietal areas exhibit differential sensitivity to rising and falling anesthetic depth. Across different anesthetic levels, we observed characteristic slow oscillations in delta range in both PFC and PPC, with neurons transitioning between high-firing “Up” states and near-silent “Down” states. Deeper anesthesia extended Down states, suppressed mean firing rates, and reduced the frequency and duration of Up states. In M1, no single units were detected in PFC, and only a few were recorded in M2. We suspect misalignment of the probe with PFC pyramidal cells and extensive suppression in PFC as the main reasons. Recurrent Neural Networks (RNN) was used to extract Up/Down states from LFP activities, based one the pattern observed in the PPC. The PPC → PFC information flow observed in Transfer Entropy analysis suggests that even under anesthesia, some level of feedforward-like interactions may persist. Up states originate in deep cortical layers and propagate toward superficial layers, following a bottom-up progression, indicating that deep-layer pyramidal neurons, which receive strong thalamic input, may be the primary drivers of Up states. The short Up states under deep anesthesia might represent a failed ignition attempt, where the brain momentarily tries to reactivate but cannot sustain functional activity due to global inhibition. LFP analyses revealed that although the absolute delta power remains high at different anesthetic levels, the relative delta band power is anti correlated with anesthetic depth, due to sporadic short (20ms to 40ms) burst in gamma (30–100 Hz) that appeared transient in nature. Lower sensitivity to anesthesia dose changes were observed in PFC as compared to PPC. This could explain why anesthesia first impairs cognitive function before affecting basic sensory responses. These results indicates that the traditional Up/Down state models might oversimplify anesthetic brain dynamics. While anesthesia is often described as a state of simple global slow-wave oscillations, the observed Up/Down state durations are not uniform, they fluctuate, follow non-trivial transition patterns, and differ between PFC and PPC.

## 1. Introduction

The posterior parietal cortex (PPC) and prefrontal cortex (PFC) are key association areas in high-level cognitive and sensory integration. In awake humans, the dorsolateral PFC and superior/posterior parietal cortex support functions such as attention, perception, and working memory (Naghavi & Nyberg, 2005). These regions are also crucial for the subjective aspects of conscious experience, as evidenced by neuroimaging studies linking PFC/PPC activity to conscious awareness (Rees et al., 2002). While some studies (Kaisti et al., 2003) observed that parietal cerebral blood flow decreases already at moderate doses of sevoflurane while frontal reductions become pronounced at higher concentrations, other studies (Hudetz, 2012) emphasize a general drop in frontoparietal metabolic activity during unconsciousness. These findings together indicate that while both PFC and PPC undergo significant suppression, the dose-response profile may differ across these regions. Even at mild sedative doses, anesthetics preferentially impair higher cognitive functions like focused attention, working memory, and executive processing (Hudetz, 2012). On the contrary, the basic sensory responses in primary cortex does not completely disappear, rather gets significantly attenuated by increase in the anesthesia depth (Kerssens et al., 2005). This means an anesthetized brain can often register incoming sensory signals in early sensory cortex, but cannot integrate them into conscious percepts or purposeful responses once PFC/PPC activity is silenced. Recent recordings in animals under propofol found that auditory and tactile inputs still activate primary sensory cortex, but these signals fail to propagate to downstream parietal and prefrontal regions under anesthesia (Tauber et al., 2024). Therefore, the “global workspace” function of frontoparietal cortex is severed: information no longer reaches the cortical areas responsible for awareness and decision-making. This breakdown in cortical communication indicates how anesthetics effectively disconnect conscious sensory processes from higher cortical areas.

While macro-scale studies point out to disruptions under anesthesia (Sorrenti et al., 2021), direct neuronal-level evidence in non-human primates (NHPs) remains comparatively sparse, especially in higher-order areas like PFC and PPC. It is not yet well understood if there are laminar differences in how these regions transition between Up/Down states under sevoflurane, and if frontal cortex truly exhibit preferential suppression? New studies are required to bridge the gap between human macro-scale data and rodent micro-scale findings, using high-density electrophysiology in NHPs. We already know that at the cellular and network level, anesthetics induce a slowing of cortical dynamics. Neurons throughout the cortex hyperpolarize under anesthesia, leading to reduced firing rates and the emergence of synchronized “slow-wave” oscillations. In both humans and animal models, unconsciousness maintained by GABAergic anesthetics (like propofol or sevoflurane) is marked on the EEG by dominant slow-delta oscillations (Adam et al., 2023). While agents like propofol and isoflurane share many features of cortical suppression, sevoflurane is more commonly used in humans and shows unique dose-dependent effects on spectral power and connectivity that needs detailed investigation (Kaisti et al., 2003; Sorrenti et al., 2021). These large slow waves reflect the cortex switching into a slow oscillation regime of neuronal spiking (Up states) alternating with silence period (Down states) of approximately once every few seconds (Torao-Angosto et al., 2021). While it is well known that anesthetics evoke global slow oscillations, it remains unclear whether local cortical layers in PFC and PPC experience uniform suppression or if certain layers preserve partial activity, a question that becomes central for understanding how different cortical depths contribute to overall unconsciousness. Moreover, evidence from our data suggests that Up states may not always propagate uniformly across PFC and PPC, raising the possibility that residual feedforward signals from parietal to frontal cortex persist even in deep anesthesia.

As anesthetic depth increases, the slow oscillation becomes more extreme and results in to burst-suppression, a pattern in which brief bursts of activity are punctuated by flatline (isoelectric) periods of suppression which is a signature of profound cortical inactivation. These slow oscillations inherently fragment normal information processing, as neurons can only communicate during the Up states and remain quiescent during Down states. This fragmented activity and significantly reduced neuronal firing both contribute to why stimuli cannot be processed under anesthesia (Adam et al., 2023). A unifying observation across studies of anesthetic-induced unconsciousness is the breakdown of effective connectivity between cortical regions. Under normal conditions, PFC and PPC form interconnected networks, knowns as the frontoparietal network, that facilitate the global integration of information. Anesthesia decouples these networks. Functional MRI studies in humans have shown that as propofol sedation deepens, functional connectivity within frontoparietal networks decreases in proportion to the loss of consciousness (Boveroux et al., 2010). This functional disconnection is a targeted disruption rather than a complete shutdown of all connections. Low-level sensory cortices often retain local connectivity and thalamic inputs even during deep anesthesia (Boveroux et al., 2010; Hudetz, 2012). These results suggest an isolated islands scenario: primary sensory areas can still talk to thalamus and to their immediate neighbors, but the long-range highways to associative cortices are effectively closed. This selective loss of long-range cortical integration is proposed to be the mechanism for the loss of consciousness and is argued that the brain under anesthesia cannot “bind” distributed information or sustain the global neuronal workspace that consciousness requires (Tauber et al., 2024). Instead, each region may process signals in a fragmentary way, preventing any unified percept or cognitive functionality.

In particular, top-down (feedback) signals from frontal cortex to posterior regions are shown to be more sensitive to anesthetic suppression. A consistent pattern across different anesthetic agents (e.g., propofol, isoflurane) is to weaken of feedback connectivity, particularly from frontal to parietal regions, while feedforward (bottom-up) sensory connectivity is relatively preserved (Boly et al., 2012). In rodents, for example, isoflurane greatly attenuates cortical responses driven by higher-order (corticocortical or “top-down” thalamo-cortical) inputs, whereas responses to direct bottom-up sensory inputs were less affected (Raz et al., 2014). These evidence all converge on the idea that anesthetics disrupt the brain’s long-distance communications and the PFC and PPC not only decrease in intrinsic activity under anesthesia; they also become functionally isolated from each other and from incoming sensory streams. However, it should be noted that most of such results are based on the propagation of sensory signals into higher cortical areas and not from analyzing the spontaneous brain activity under anesthesia, and particularly the up and down states interplay between distant areas.

While the core phenomena of anesthesia – unconsciousness, slow oscillations, connectivity loss – are broadly observed across mammals, there are important variations between species and even between individuals. Across species, primates appear to retain more localized cortical activity during burst-suppression compared to rodents. In rats the entire neocortex tends to enter synchronous suppression, whereas in primates many primary sensory areas continue to show activity during bursts (Sirmpilatze et al., 2022). Primate sensory areas are heavily interconnected making them more resistant to global shutdown. Within a species variability in anesthetic response is also notable. Both in humans and NHPs, while general anesthesia significantly reduces inter-individual differences in brain connectivity, it does not entirely eliminate inter-individual variability, with the highest variability remaining in higher-order cognitive regions such as the default mode network. In contrast, sensory and motor regions, which are more structurally constrained, show the least variability under anesthesia (Luppi et al., 2023; Xu et al., 2019). Factors like age, physiological state, or genetics can explain how the interindividual variability and how different cortical areas of a given animal react to the drug. Understanding why one there is relatively large inter-individual variance withing species is crucial for advancing personalized anesthesia protocols and bridging important translational gaps to clinical practice. These findings indicate that while the overall effects of anesthesia is shared between and across species, the precise spatiotemporal pattern can vary significantly across species, and depending and the cortical area and recording scale withing species. The challenge is that the fine details of neuronal dynamics under anesthesia can look dissimilar in smaller brains versus larger brains. This reinforces the importance of studying both local circuits and global functional connectivity.

While macroscopic effects of anesthesia are widely studied, the literature on the microscopic effect and intra regional neural pattern is scarce, particularly in larger models and NHPs (Sorrenti et al., 2021). Laminar recordings provide a depth-resolved view of cortical columns, revealing, for instance, that deep anesthesia strongly suppresses supragranular activity (associated with feedback inputs), and less on the thalamocortical input to granular layers. These laminar information can help to explain EEG phenomena like the amplification of low-frequency waves – they originate from synchronized dendritic currents during cortical Down states – and the diminution of higher frequencies – due to reduced local recurrent firing. A study on ferrets reported that isoflurane caused distinct laminar effects. In V1, it enhanced LFP spectral power predominantly in granular (IV) and infragranular (V-VI) layers and increased visually evoked firing in layer IV. In contrast, in PFC, the modulation observed in layers II/III during visual stimulation in awake animals was lost under anesthesia (Sellers et al., 2013). Using the information-based theory on the same dataset, it has been suggested that isoflurane anesthesia induces layer- and frequency-specific changes in cortical information processing. In supergranular layers (upper layers I-III, involved in inter-cortical communication), modulation predominantly occurs in the high/low gamma band. In infragranular layers (deeper layers V-VI, responsible for outputs to subcortical regions), the most significant decrease in information retention is observed in the alpha/beta band (Pinzuti et al., 2023). Another study using laminar recording in frontoparietal loop while stimulating thalamus highlights the critical role of layer-specific neural interactions in consciousness. It found that deep cortical layers, in collaboration with the central lateral thalamus, sustain activity essential for conscious states and argues that feedback from deep to superficial cortical layers plays a key role, emphasizing the importance of long-range feedback and intracolumnar signaling in consciousness regulation (Redinbaugh et al., 2020). Another study in rats demonstrates the initiation of slow wave activity (SWA) in the somatosensory cortex under ketamine anesthesia is largely driven by firing in layer V, while layer IV also plays a role, particularly under somatosensory stimulation. This study highlights the spatially distinct roles of cortical layers in SWA generation, emphasizing the involvement of thalamic input and local cortical dynamics (Fiáth et al., 2016). It is important to note that these findings are under ketamine, which has a different mechanism and impact compared to sevoflurane and isoflurane, as discussed in earlier studies. Collectively, there is a consistent observation that anesthesia affects supragranular vs. infragranular layers differently. High-dose barbiturate anesthesia, for example, suppressed most spiking and LFP responses in superficial layers, even while deeper layers continued to exhibit activity. Deep-layer neurons often remain active or bursty despite anesthesia. Layer V pyramidal cells can continue to generate bursts and even initiate Up states when superficial layers are quiescent (Kuo et al., 2015). It has been suggested that deep cortical layers have mechanistical role in consciousness level under anesthesia (Hönigsperger et al., 2024). Laminar recordings also allow researchers to investigate layer-specific feedback/feedforward disassociations under anesthesia. For instance, findings from multi-site laminar recordings in rats shows how feedback from higher-order cortex to layer 5 in V1 is completely lost under anesthesia (Hudetz et al., 2020), reinforcing the idea that anesthesia functionally disconnects cortical circuits, both locally (between layers) and globally (between regions), indicating how anesthetic-induced unconsciousness involves a breakdown in both horizontal and vertical integration of cortical processing. Importantly, this breakdown is most pronounced in association cortices like the PFC and PPC, which normally rely on rich interlaminar communication for higher cognitive functions. Under anesthesia, frontal and parietal regions exhibit strong low-frequency synchrony but diminished information flow, essentially entering a local “default” oscillatory mode decoupled from broad network interaction. This is consistent with human studies pointing to frontoparietal disconnection as a hallmark of anesthetic unconsciousness. Examining the laminar patterns in PFC and PPC under anesthesia (as in the present study) is critical – these regions show the clearest signatures of layer-specific suppression and disrupted connectivity, linking the microcircuit effects of anesthesia to the macroscopic loss of consciousness.

Collectively, the existing literature shows that although anesthesia suppresses conscious processing, its deeper mechanisms, ranging from laminar activity gradients to global and local patterns of Up/Down states, remain only partially understood. Our preliminary observations in macaque PPC and PFC, which reveal notable differences across in laminar firing-rate pattern and between individuals, highlight the need for a detailed investigation that can clarify how these circuits respond to sevoflurane-induced inactivation. Understanding whether deeper layers of cortex are driving or receiving inputs during Up-state transitions, how these transitions compare in the different cortical areas, and whether Up/Down states are globally synchronized or arise from local, recurrent processes is crucial for clarifying the interplay between slow-wave synchronization and residual network dynamics. With advances in high-density recording technology, it is now possible to examine these questions at a finer scale in larger animal models. Using Neuropixel probes in the PFC and PPC of two macaque monkeys, we aim to fill these knowledge gaps at a level of detail previously not feasible. Such findings are essential for advancing theories of top-down vs. bottom-up cortical disintegration under anesthesia and for tailoring anesthetic strategies to each subject’s unique cortical reactivity. In doing so, we not only test established ideas, such as the primacy of frontoparietal disintegration in the loss of consciousness, but also provide new insights into how layer-specific dynamics, transient gamma bursts, and feedforward influences shape the anesthetized brain.

## 2. Materials and Methods

### 2.1 Subjects and Ethical Approval

Two adult male macaque monkeys (M1, 21 years old, 14 kg; M2, 18 years old, 16 kg) were used in this study. Both were pair-housed under a 12-hour light/dark cycle with free access to food and water. All procedures were conducted in accordance with national guidelines for the care and use of nonhuman primates and approved by the institutional animal care and the internal committee. The animals had previously undergone separate surgeries to implant Utah arrays in the posterior parietal cortex and the dorsolateral prefrontal cortex bilaterally.

### 2.2 Surgical Preparation and Probe Implantation

Monkeys were initially sedated with ketamine (approximately 10 mg/kg i.m.), supplemented by a standard opioid analgesic (e.g., 0.01 mg/kg buprenorphine i.m.) before transport to the operating room. Sevoflurane inhalation anesthesia was then introduced and maintained via endotracheal tube. The scalp was shaved and disinfected with povidone–iodine, after which a midline incision exposed the skull and allowed the periosteum to be carefully removed. As indicated by the landmarks from the prior implants, a small (∼2 mm) craniotomy was performed above each cortical targets in PPC and PFC (Figure 1B). The dura was lightly scratched using the surgical needle before placing the Neuropixel 1 probes (10 mm shank, configured for 384 active recording channels along its length). Since the entire preparation and implantation lasted more than two hours, any residual ketamine effects had sufficiently washed out before the start of recordings.

**Figure 1.**
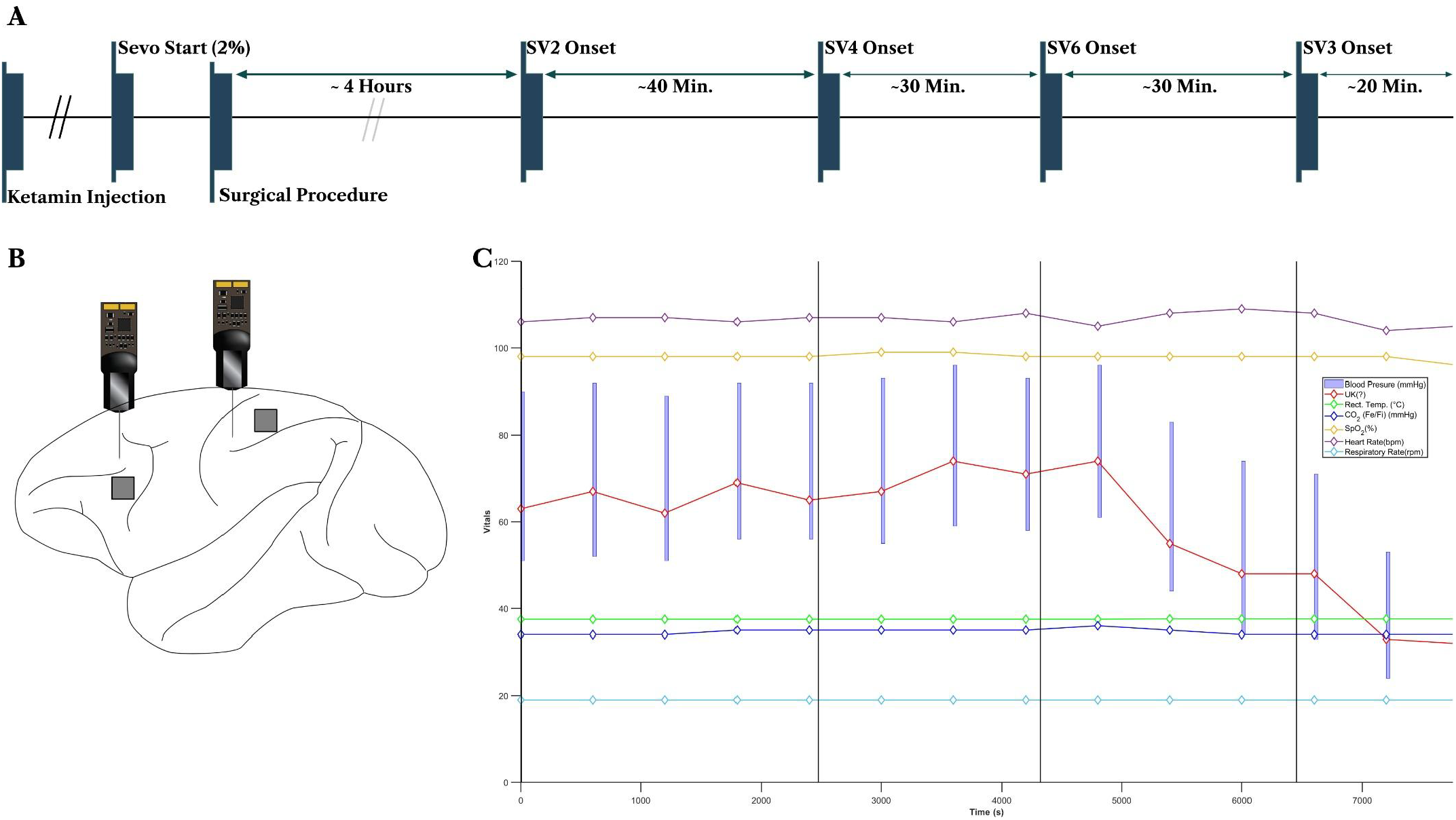
Overview of the anesthesia protocol and physiological monitoring. (A) The experiment proceeded through four sevoflurane levels (2%, 4%, 6%, and 3%), each maintained long enough for stable recordings. (B) Vitals remained largely within safe ranges, Blood pressure (systolic, diastolic) and heart rate with fluctuationed shortly after the transitions and anticorrelated with the anesthesia depth. (C) Putative locations of existing Utah arrays and newly implanted Neuropixel probes in both PPC and PFC, as inferred from surgical landmarks during Utah implantation.

### 2.3 Experimental Design

Four sevoflurane levels were administered consecutively: 2% for about 40 minutes (SV2), followed by 4% (SV4) for 30 minutes, 6% (SV6) for 20 minutes, and finally 3% (SV3) for 20 minutes. Transitions between levels occurred after vital signs (e.g., blood pressure, respiratory rate, SpO_2_, end-tidal CO_2_) had reached stable values. Figure 1A provides an overview of the experimental design, including the sequential anesthesia protocol and approximate time spans at each concentration. Figure 1C illustrates representative physiological recordings confirming stable anesthetic depth.

### 2.4 Data Acquisition and Preprocessing

Neuropixel signals from both PPC and PFC were acquired using OpenEphys (Siegle et al., 2017). Spikes were sampled at 30 kHz; LFPs were recorded at 2.5 kHz. Using Spikeinterface (Buccino et al., 2020), the raw spike data were high-pass filtered between 600 Hz to 12.5 kHz, corrected for phase shifts, and referenced using common average reference. Automatic detection flagged bad channels, which were then visually inspected; no major motion artifacts or drift were observed using spikes’ peak locations. Three to five dominant noise frequencies were removed using narrow width notch filters. Spike sorting was conducted with Kilosort 4 (Pachitariu et al., 2024) and curated in multiple steps using Spikeinterface curation module. Units were initially labeled by firing rate, signal-to-noise ratio, and refractory violations. Next, manual inspection merged or removed questionable units based on amplitude distributions, autocorrelograms, template metrics and template shapes (Jung et al., 2023). A total of 234 well-isolated single units in PPC of M1, and 204 in M2 passed final inclusion criteria along the shank. Although LFP signals were robust in both cortical regions, spiking activity in PFC was largely absent (see Section 2.5). LFPs were filtered between 0.5 Hz to 90 Hz. Instantaneous band power were extracted from Hilbert transform of the filtered signal in their respective band frequency range (Berens et al., 2008). All subsequent power spectral analyses and inter-regional connectivity measures were carried out in MATLAB 2023 and Python 11.4 using standard toolboxes (Oostenveld et al., 2011). Wherever multiple frequency bands or time windows were compared, Bonferroni corrections were applied to control the familywise error rate, and significance was set at α = 0. 05.

### 2.5 PFC Single-Unit Yield

Despite robust LFP signals in PFC, high-quality single units were rarely obtained. The poor template metrics and high noise levels for putative units suggested either severe anesthesia-related suppression or unfavorable neuron–electrode orientation. In M2, a handful of spikes in deeper channels did pass curation thresholds for good signal-to-noise ratios and consistent waveforms. For this monkey, we used a relatively high threshold (15 MAD, median average deviation) as for detecting the multi unit activities (MUA) for getting the population activity and the up and down states. Overall, the inability to isolate well-defined PFC units in the face of normal LFP signals suggests a predominantly physiological (rather than technical) phenomenon.

### 2.6 Neural Data Analysis and Up/Down State Classification

#### 2.6.1 PPC Up and Down State Detection

Three complementary methods were applied to spiking activity in PPC to classify cortical Up and Down states:

1. **Moving Average Thresholding:** We first convolved the spike train with a rectangular window of length *w* to obtain local mean 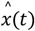. Next, we computed a local standard deviation over the same window. At each time bin, an Up state was assigned if the instantaneous spike count *x*(*t*) exceeded 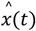 plus n times the local standard deviation. Formally,

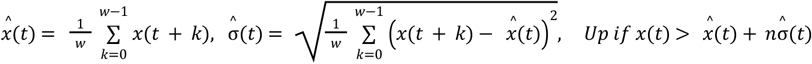 A final correction step merged isolated single-bin Down states bounded by Up states into an Up state to avoid brief, spurious fluctuations.
2. **Phase-Space Reconstruction:** The spiking time series *x*(*t*) embedded into a 3-dimensional phase space using an embedding delay *τ*. For each time *t*, we constructed

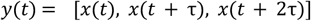 Distinct clusters in this phase space were identified as Up versus Down states by minimizing the within-cluster sum of squares (MiniBatch K-Means algorithm):

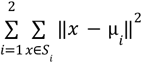 Where *S*_*i*_ is the set of point in cluster *i* and μ_*i*_ is the centroid of cluster *i*. Up states corresponding to regions of high spike density or rapidly changing population activity.
3. **Hidden Markov Model (HMM):** We first applied a light Gaussian filter (σ = 1) to the binned spike activity, then standardized the smoothed signal. A Gaussian HMM with four latent components was fit to the resulting time series, allowing the model to capture different amplitude regimes of spiking activity. After training, each time bin was assigned to a hidden state. The state exhibiting the largest emission mean was relabeled as “Up”, while the remaining states were considered “Down”. A brief gap-correction step merged single-bin Down interruptions bounded by Up bins into the Up state to reduce spurious fluctuations.

After applying all three methods across different sevoflurane levels, we visually inspected how closely the detected Up and Down states aligned with population spiking activity. The moving-average approach resulted in frequent false positives during prolonged Down states under deeper anesthesia. Phase-space reconstruction performed better, but still was contaminated by spurious fluctuations specially in lighter anesthesia with frequent up states, resulting in shorter, more frequent Up states. By comparison, the HMM was more stable and consistent with observed transitions across all anesthesia depths. Although no independent ground truth was available, these side-by-side evaluations at different anesthetic levels strongly suggested that the HMM provided the most accurate characterization of the underlying neural dynamics.

#### 2.6.2 PFC Up/Down State Estimation via LFP Modeling

Because the PFC yielded minimal spiking activity, particularly in M1, a recurrent neural network (RNN) was trained to learn Up and Down state labels from PPC LFP signals alone and then applied to PFC LFP data. Principal components (PCs) explaining 95% of the variance in the PPC LFP were extracted and labeled according to the spike-based HMM-derived Up/Down states. These labeled PCs fed a bidirectional LSTM network (64 units, 20% dropout, followed by 32 units, 20% dropout, and a sigmoid output). To address the large imbalance between Down and Up states (∼24:1), a focal loss function was hired,

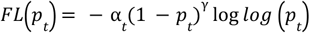

where *p*_*t*_ is the predicted probability of the true class, α is the weight for balancing the classes, and γ is the focus parameter to downweigh easy examples. Class-specific precision, recall, and F1 scores guided hyperparameter tuning, and the trained model was then used to infer Up/Down states in the PFC solely from its LFP principal components. For M2, we used the MUA to detect up and down states, with similar method explained in 2.6.1.

### 2.7 Transfer Entropy Analysis

Transfer entropy (TE) was computed in a sliding window of 30 seconds with 0.5-second overlaps, using the open-source TRENTOOL3 toolbox (Lindner et al., 2011). TE estimation followed the estimator guaranteeing optimal self prediction method (Wibral et al., 2013), scanning a range of interaction delays from 50 to 500 ms in 10 ms increments. Trial-based shuffling with 50,000 permutations tested for statistical significance (one-tailed, with TEshift>TE). TE was then calculated bidirectionally (PPC→PFC and PFC→PPC) to detect potential asymmetric information flow, as well as the diagonal cases (PPC→PPC and PFC→PFC) for baseline comparison. The output included estimates of statistically significant directional influences in each 30-second segment across the full recording, indicating a time-resolved measure of how interactions changed with varying sevoflurane concentrations.

## 3. Results

### 3.1 Behavioral and Physiological Monitoring

Throughout the experimental sessions, both monkeys remained in a stable physiological range. Heart rate and blood pressure measurements (Figure 1C) varied in response to changes in sevoflurane concentration, with brief, transient surges typically observed immediately after each new dose was introduced. However, these fluctuations did not exceed typical sedation ranges and quickly returned to a stable baseline, suggesting that the dose transitions (2% → 4% → 6% → 3%) were well-tolerated overall. No abrupt or sustained deviations from normal cardiovascular parameters were noted.

### 3.2 LFP Spectral Reorganization

Power spectral density (PSD) analyses and time-resolved band power measurements revealed a linear decline in relative delta-band (0.5–4 Hz) power from approximately 51% to 34% of total power as sevoflurane concentration increased (SV2→SV3→SV4→SV6), in contrast to more conventional reports of enhanced delta under anesthesia (Figure 4A). By comparison, gamma-band (30–90 Hz) power remained at roughly 5% across SV2–SV4, then rose to about 35% at SV6. A similar pattern was observed for theta (4–8 Hz), which showed a linear increase up to SV4 (20% to 30%) followed by a sudden drop to 10% at SV6. It is important to note that these values refer to relative power: when one band (e.g., gamma) increases disproportionately, other bands’ percentages appear lower even if their absolute values remain unchanged. This effect is further amplified by the fact that gamma spans 60 Hz (30–90) while delta covers only 4 Hz.

Both alpha and beta remained stable across SV2–SV4, with alpha decreasing from ∼10% to 8% of total power at SV6, whereas beta rose from ∼12% to 15%. Despite the overall increase in relative gamma at deeper anesthesia, discrete gamma surges also emerged. These bursts started ramping up shortly after transitioning to SV6, suggesting that abrupt changes in anesthetic depth trigger transient network reconfiguration (Figure 4C). Within ∼10–15 minutes of reaching SV6, burst frequency stabilized, though it remained higher than at lighter anesthetic levels. A similar surge also occurred when moving from SV6 back to SV3, again presumably reflecting the cortex adapting to a new sedation level. Notably, the emergence of these gamma bursts does not correlate with an increase in the number or duration of Up states, and their functional significance, whether they signify partial network re-engagement or localized excitatory events, remains unclear.

These observations held for both PPC and PFC. However, in PFC the gamma bursts were less frequent and took longer to stabilize after changes in anesthesia. PPC showed higher variance in gamma power across channels than either other bands or PFC. In PPC, deeper layers showed relatively higher delta, while superficial layers exhibited stronger gamma. Furthermore, certain channels that already exhibit elevated gamma power, increased further under deeper anesthesia, extending to neighboring channels. By contrast, PFC showed less pronounced laminar clustering of gamma, with more power consistently concentrated in the delta range and only a modest gamma increase at SV6 (Figures 4C). Analyzing the 1/f slope in PSDs using the FOOOF method showed that, in PPC, both the offset and exponent parameters clearly separated the four anesthesia levels. While in PFC, SV2, SV3, and SV4 largely clustered together and diverged only at SV6 (Figure 4B). Overall, while both areas shifted toward higher gamma and lower delta under deeper anesthesia, PPC responded more robustly and more quickly, whereas PFC changed more gradually and displayed a pronounced spectral shift only at the highest concentration.

Across both PPC and PFC, relative band power fluctuations revealed prominent negative correlations among frequency bands. Delta and beta were most strongly anti-correlated (PPC: r = –0.56; PFC: r = –0.54), with similar inverse relationships between delta and both theta and alpha, suggesting that increases in slow oscillatory activity tend to suppress neighboring frequencies. In PPC, delta and gamma were also negatively correlated, likely reflecting transient gamma bursts that draw power away from lower frequencies. In contrast, beta and gamma showed modest positive correlations (PPC: r = 0.33; PFC: r = 0.18), indicating that mid- and high-frequency activity can sometimes co-vary. It is important to note that these data were assessed as relative band power, meaning the sum of all bands at each time point is normalized to 1. By definition, a rise in one band causes a proportional drop in another. Taken together, the observed patterns suggest a dynamic push-pull between slow and fast rhythms under anesthesia, with brief shifts into beta–gamma coactivation potentially marking transient desynchronization or “micro-arousal” states within an otherwise slow-dominated regime.

### 3.3 Overview of Spiking Activity Across Anesthesia Levels

Neuropixel recordings in the posterior parietal cortex (PPC) identified 253 well-isolated units in Monkey 1 (M1) and 275 in Monkey 2 (M2). By contrast, no consistently isolable units were detected in M1’s prefrontal cortex (PFC), suggesting pronounced physiological suppression rather than a technical artifact. M2’s PFC yielded 10 stable units, all confined to deeper channels, yet their overall firing rates remained low.

As sevoflurane concentration increased, spiking in PPC progressively declined to near-zero during extended silent intervals up to tens of seconds, highlighting a profound suppression of cortical output at deeper anesthetic levels (Figure 2A). Transitions from lower to higher anesthesia levels (2% to 4% and 4% to 6%) require approximately 10–15 minutes to reach a stable firing pattern, whereas transitions from higher to lower anesthesia levels (6% to 3%) stabilize more rapidly, within 3–5 minutes. These temporal dynamics closely align with patterns observed in LFP recordings, though with a longer timescale, possibly reflecting the longer neuronal or synaptic recovery under anesthesia.

**Figure 2.**
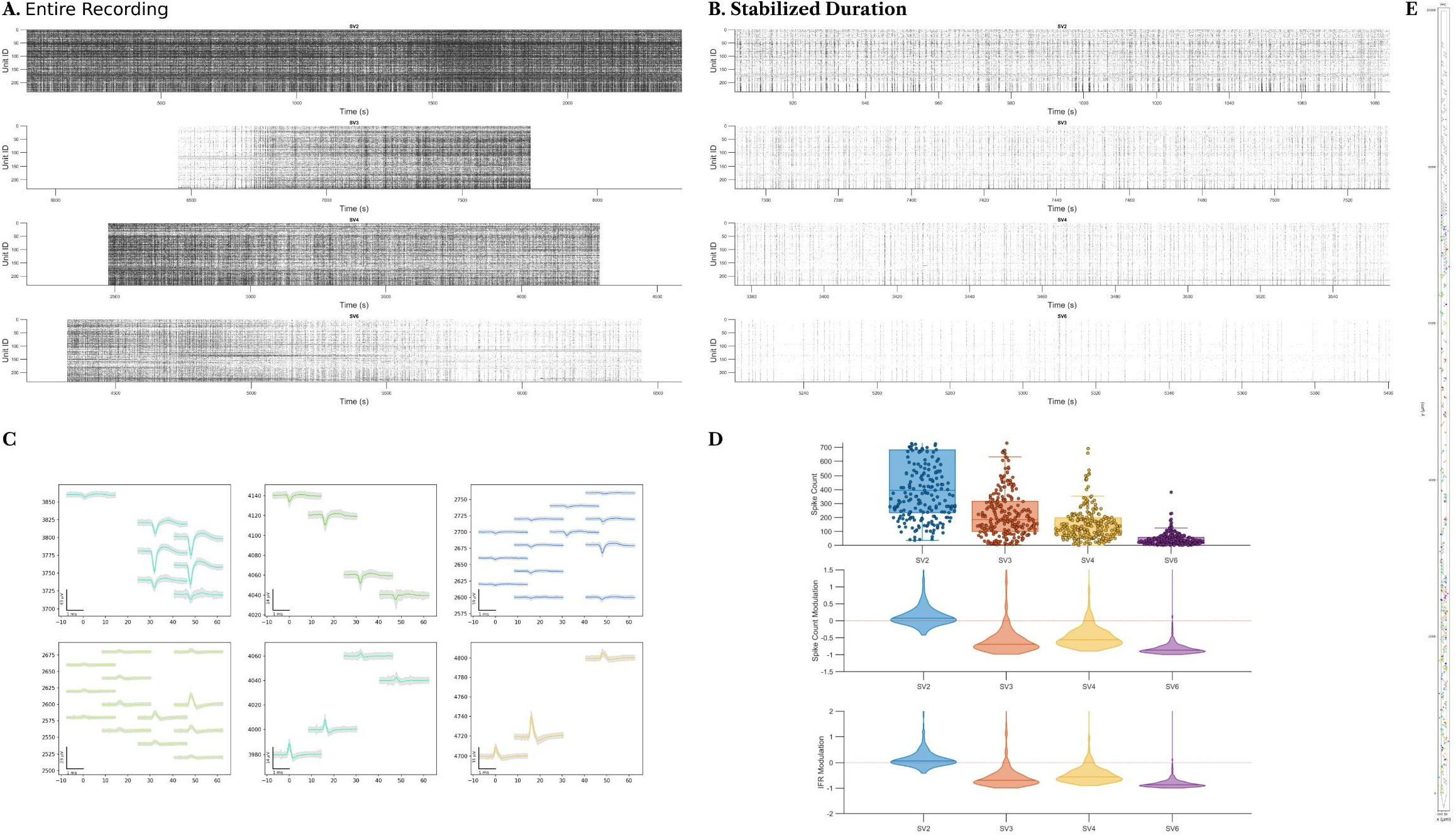
Spiking profiles under anesthesia (M1). (A) A raster plot of the entire recording session for one anesthesia level captures the global suppression and occasional bursts. Subplots are sorted based on the anesthesia depth, while 3% sevoflurane came after 6%, hence the gradient recovery. (B) A zoomed-in raster during the stabilized phase (3 minutes, beginning 15 minutes after an anesthetic shift) reveals clearer Up/Down fluctuations. (C) Handpicked single-unit waveforms in PPC illustrate the variety of spike shapes isolated. Same unit is recorded with different channels. (D) Overall spike count (Top), and spike count and average Instantaneous Firing Rate (IFR) modulation (Middle and Bottom, respectively) relative to the initial 5 minutes of the recording (baseline), demonstrating progressive suppression under deeper anesthetic concentrations. (E) Spatial organization of the detected units along the shank for PPC.

Using the first five minutes of SV2 as baseline, we evaluated spike count and instantaneous firing rate modulations during the stabilized period (15 minutes post-dose change over a 3-minute window) for each condition. Both metrics declined linearly with escalating anesthetic dose, culminating in up to 100% suppression of at the highest levels (Figure 2D).

After anesthesia stabilization, down-state durations grew longer with each incremental increase in anesthesia (Figure 2), indicating prolonged neuronal inactivity at deeper sedation. At lighter concentrations, the network alternated between up and down states more frequently, whereas at SV6, long and irregular down states dominated. Collectively, these findings points out the strong dose-dependent impact of sevoflurane on spiking dynamics in PPC and PFC, including marked differences between the two monkeys (See 3.4) and an apparent frontoparietal gradient in anesthetic-induced neuronal suppression.

### 3.4 Up and Down State Dynamics

Under all anesthetic levels, PPC exhibited pronounced Up/Down oscillations, characterized by brief epochs of coordinated firing (Up states) interspersed with prolonged silences (Down states) (Figures 2B, Figure 3A). To detect these states from population spike activity in PPC, we implemented three methods, Hidden Markov Model (HMM), Phase Space Reconstruction, and a Moving Average approach. Visual inspection revealed that HMM most reliably captured transitions as the Phase Space Reconstruction introduced sporadic state changes, and the Moving Average approach produced excessive false positives.

**Figure 3.**
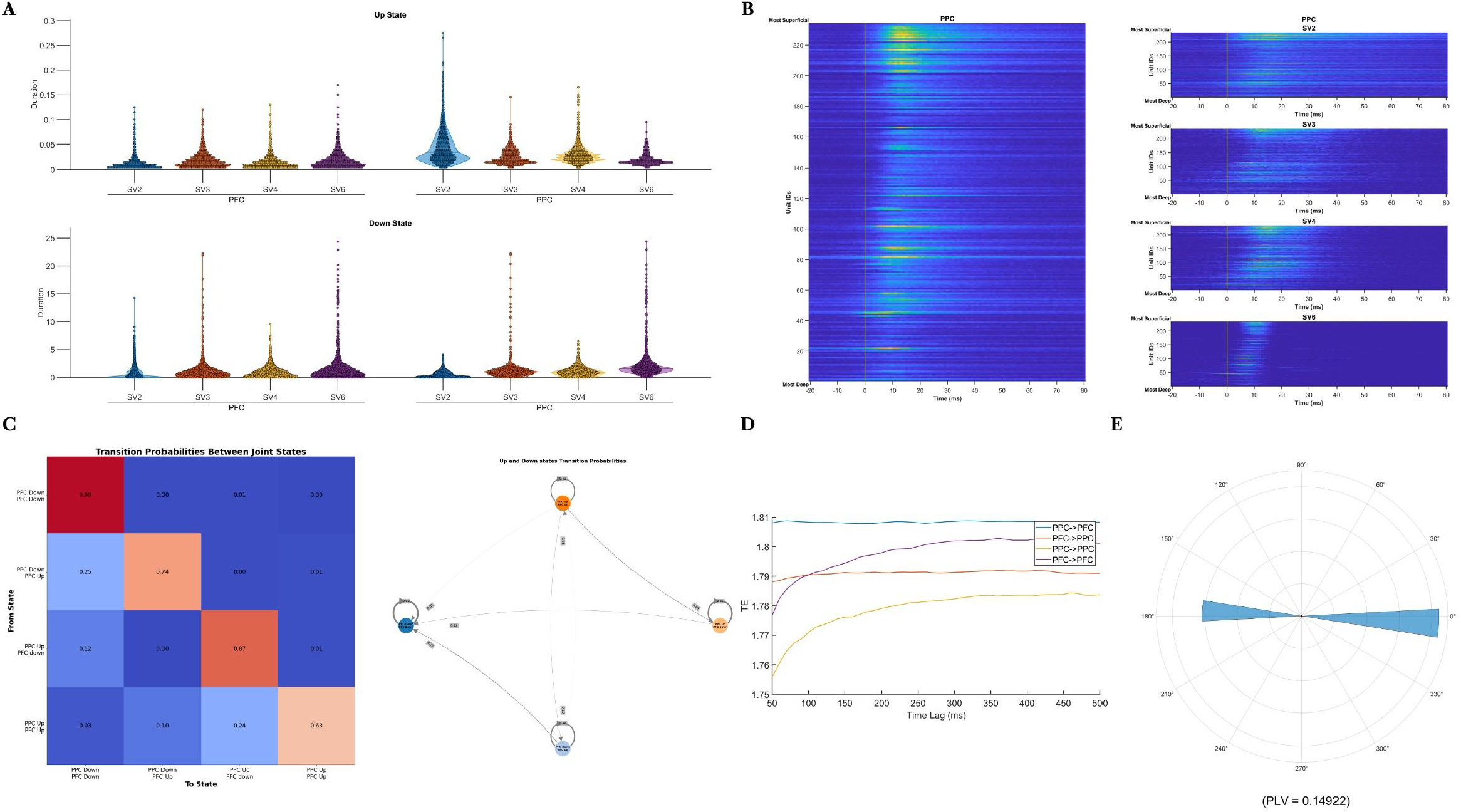
Up/Down state dynamics under sevoflurane (M1). (A) Violin plots of Up and Down state durations separately for PPC and PFC. (B) Spatiotemporal progression of Up states in PPC indicates that neuronal firing typically initiates in deeper layers and propagates superficially, capturing the laminar wave of activity during each Up event. (C) Matrix representations (Left) and Graph-based (Right) of Up/Down transition probabilities. (D) Transfer entropy (TE) analysis how well each area predicts itself and the other over different time lags. (E) Phase lag difference between PPC and PFC Up and Down state transitions.

**Figure 4.**
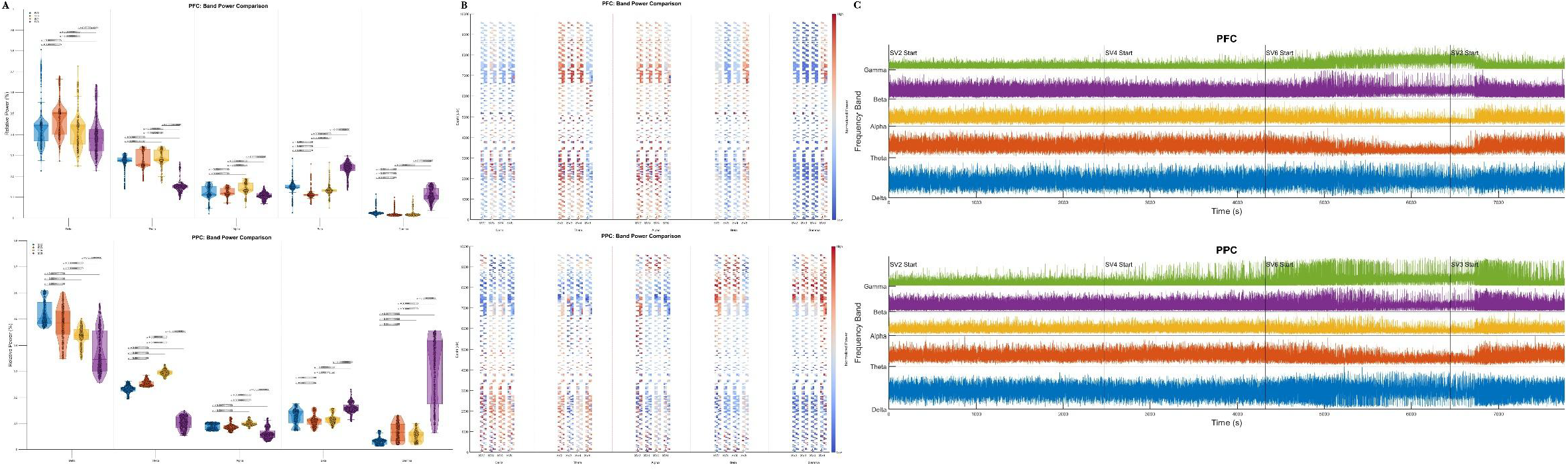
LFP power reorganization and instantaneous band dynamics. (A) Grouped comparisons of relative band power for PFC (Top) and PPC (Bottom). (B) Laminar distribution power the relative power across different channels along the shank for PFC (Top) and PPC (Bottom). The heatmap color is normalized for each channel individually (and not globally). (C) Traces of instantaneous band power, computed via Hilbert transform depicting moment-to-moment fluctuations for PFC (Top) and PPC (Bottom).

We next trained a long short-term memory (LSTM) model to classify Up/Down states using both spike and LFP signals in PPC, to be able to infer the up and down states in PFC only from LFP activity, achieving an overall classification accuracy of 98%. Given the 24:1 up-to-down state imbalance, we prioritized class-specific metrics, particularly precision and recall, to evaluate the model’s performance. Analysis of the precision-recall curve (Supplementary Figure XX) showed a trade-off in which higher recall for Up states came at the cost of losing precision, and vice versa. To optimize the precision-recall balance, we selected a threshold of 0.4, which resulted in F1-score of 0.99 for Down states and F1-score = 0.76 (Precision = 0.72, Recall = 0.80) for Up states. This threshold provides a balanced model performance, prioritizing accurate detection of down states while maintaining reasonable sensitivity for up states.

With increasing anesthesia depth, down-state duration progressively increases, with the mean duration extending from 0.9 seconds to approximately 3 seconds, reaching up to 25 seconds after stabilization in M1. This pattern is consistent between PFC and PPC. Up-state durations in PPC decrease from an average of 25 ms (SV2) to 15 ms (SV6), indicating a progressive shortening of active periods as anesthesia deepens. In PFC, however, up-state duration increases from 10 ms (SV2) to 25 ms (SV6), suggesting a region-specific effect of anesthesia on cortical state transitions. The observed changes in both up-state and down-state durations follow a linear progression across anesthesia levels (SV2 → SV3 → SV4 → SV6), indicating a gradual and systematic modulation of cortical dynamics with increasing anesthetic depth. In M2’s PPC showed the longest Down states at 6% anesthesia, often lasting over a minute in certain segments. Figure 3A illustrates the distribution of Up and Down state durations across conditions.

Layer-specific analysis reveals that up-states originate in the deepest cortical layers and progressively propagate toward the more superficial layers (Figure 3B). This bottom-up progression of up-states is particularly pronounced in the deepest anesthetic state, where background neuronal activity is significantly reduced and neural synchrony is higher, making the sequential activation pattern more distinct. Since there are no histological data available, we are not sure which do these lower channels are located within the cortical layers. However, as the deep-layer pyramidal neurons, especially in Layer V, have strong recurrent excitatory connections and are more likely to initiate bursts of activity that drive up-state transitions we speculate these electrodes are located in layer V. Layer V also has dense thalamocortical input, which can contribute to the generation of up-states, particularly under anesthesia.

Cross-correlation analysis between up and down states in PPC and PFC reveals no evident correlation at different lags from −500ms to +500ms. At 10ms lag, the average correlation increases to 0.06 from 0.02, suggesting a slight temporal offset in state transitions between these two regions. Accordingly, the Phase Lag Value (PLV) analysis indicates minimal synchrony between up and down states in PPC and PFC (PLV = 1.5). However, the two signals predominantly exhibit either in-phase or anti-phase alignment, meaning they tend to be either synchronized or completely out of phase, rather than showing intermediate phase relationships (Figure 3E).

Transition probability analysis between the four cortical states (PPC-Up, PPC-Down, PFC-Up, PFC-Down) reveals a strong state persistence effect, meaning that once a region enters an up or down state, it tends to remain in that state for an extended period. The most significant transition pattern is observed when PPC is in a down state and PFC is in an up state in this scenario, the PFC has a higher probability of transitioning to a down state shortly after. Additionally, when both PPC and PFC are in an up state, the PFC tends to transition to a down state earlier than PPC, consistent with the previously observed shorter PFC up-state durations compared to PPC (Figure 3C). This suggests that PFC up-states are more transient, while PPC maintains a more sustained up-state activity. Transfer Entropy (TE) analysis over 3-minute windows across the entire recording period reveals that PPC systematically predicts PFC activity across various time lags (50 ms to 500 ms) at a higher magnitude than any other causal interaction (e.g., PPC → PPC, PFC → PPC, or PFC → PFC). Self-prediction (PPC → PPC and PFC → PFC) exhibits the lowest entropy values, particularly at shorter time lags, suggesting that local cortical dynamics are less informative about their own future states compared to cross-region influences. The optimal time lag for information transfer between PPC and PFC peaks ∼200ms, indicating that PPC activity most effectively predicts PFC activity at this delay (Figure 3D).

### 3.5 Subject-Specific Variability (M1 vs. M2)

Both M1 and M2 exhibited the main spectral and spiking trends under escalating sevoflurane concentrations, but they differed considerably in how intermediate anesthetic levels (3% and 4%) clustered. In M1, these concentrations resembled the lighter 2% condition with respect to relative band power and up-state frequency. By contrast, M2’s 3% and 4% conditions aligned more closely with its deepest (6%) state, suggesting an overall deeper sedation profile. Indeed, M2 frequently showed down states lasting over a minute at 6%. This discrepancy likely reflects multiple factors (age, metabolic rate, or subtle methodological variations) highlighting the importance of individualized anesthetic calibration. Additionally, the shorter (14-minute) recording at 3% in M2 may have prevented complete stabilization, thereby obscuring any clear rebound of spike activity following 6%.

Laminar segregation of delta and gamma in M2’s PPC was also less pronounced than in M1. Nonetheless, M2 did present a small set of large-amplitude (≥15 MAD) PFC units that enabled direct Up/Down state extraction. Applying the same LSTM-based classification parameters used for M1 achieved 95% accuracy in M2’s PFC; however, the precision–recall balance remained suboptimal, with the F1-score for Up states not exceeding 0.6. This suggests that detecting Up states in M2’s PFC was more challenging, possibly due to lower baseline spiking or reduced stabilization. Collectively, these findings underscore the significant subject-specific variability in cortical network responses to sevoflurane and emphasize the need for tailored analytical approaches (Supplementary Figures XX to YY).

It is also important to note that we set the sevoflurane vaporizer to specified percentages without measuring end-tidal (expired) concentrations. Hence, factors such as respiratory efficiency, pulmonary function, or individual metabolic rates could have led to different effective doses at the cortical level despite apparently identical vaporizer settings. In other words, one subject’s 3% might functionally be another’s 4% or 6%, leading to deeper sedation in some cases and contributing to the discrepancies observed between M1 and M2.

## 4. Discussion

In this study, we recorded laminar neural activity from the PPC and PFC of two macaque monkeys at four sevoflurane concentrations (2%, 3%, 4%, and 6%). Despite the robust LFPs in both areas, the spiking activity in PFC was either nearly absent (M1) or severely sparsed (M2). Spectral analyses of LFPs showed a systematic decrease in relative delta-band power and an increase in gamma at higher anesthetic levels in both PPC and PFC, while absolute delta-band still occupied the largest share of total power under all anesthesia levels. The pronounced low-frequency oscillations observed across PFC and PPC are in inline with reports of emergence of slow-wave state under unconsciousness (Adam et al., 2023; Steriade et al., 1993). However, the relative decline in delta band power and a striking increase in gamma at deeper anesthesia is contrary to classical reports of delta enhancement under unconsciousness. This discrepancy is partially amplified due to the focuse on relative (rather than absolute) power, meaning that large proportional rises in gamma overshadow the delta band percentage even if delta itself remains substantial. In other words, the surge in gamma power “pulls down” the relative share of lower frequencies. Spontaneous band power marked an increase in transient gamma bursts at deeper anesthetic levels, challenging the notion that only slow-wave activity dominates in profound sedation. Transient rises in gamma under anesthesia have been reported in other studies, where they may mark short-lived local cortical reactivations that fail to achieve global integration (Hwang et al., 2013; Lord et al., 2023). Our data suggest that these bursts appear more frequently at higher doses of sevoflurane, especially in the PPC, probably because the bottom-up pathways can retain partial functionality even as top-down (frontal) networks shut down. The present findings thus resonate with the broader consensus that frontoparietal disconnection, rather than total cortical silence as speculated by the wet blanket hypothesis (Voss & Sleigh, 2021), is pivotal in anesthetic-induced unconsciousness. Although both monkeys generally followed the same broad trends, substantial inter-individual variability emerged at intermediate anesthetic levels, suggesting that middle range anesthetics can differently cluster with light or deep anesthesia depending on subject-specific physiology.

The absence of well-isolated single units activity in PFC may reflect anesthesia’s known tendency to strongly suppress frontal cortex (Bastos et al., 2021) and sevoflurane role in suppressing medial PFC (Hudetz, 2012), but it could also arise from the PFC’s morphology which leads to suboptimal probe alignment. Extensive gyrification of the macaque’s PFC makes it difficult to achieve perpendicular penetrations with linear electrode arrays (Dotson et al., 2024). Macaque’s PFC is partially buried within the principal sulcus, further complicating access for laminar electrophysiology. To address the absence of direct spiking measures in PFC, we trained a deep neural network model on PPC’s spike–LFP relationships to infer analogous up and down states in PFC. The analysis revealed well-defined up/down state oscillations in PPC at all sevoflurane doses, with deeper anesthesia producing longer Down states and shorter Up states. Inferred up/down states in PFC followed broadly the same pattern, though PFC’s Up states were typically briefer, consistent with a frontally biased suppression gradient. These findings confirms that sevoflurane anesthesia induces global slow oscillations across frontal and parietal regions while also highlighting regional differences in spiking responsiveness, laminar activation profiles, and susceptibility to dose-dependent shifts in oscillatory power. This goes in line with finding of a recent study on the macaque brain which indicated that propofol anesthesia consistently destabilizes neural dynamics across cortical areas, while the degree of destabilization varies significantly between regions, with the PFC exhibiting relatively lower destabilization compared to the PPC, making the PFC less responsive to perturbations (Eisen et al., 2024).

Quantified inter-regional state transition probabilities indicated minimal synchronization between the two regions’ at lags up to ±500 ms. This suggests that PPC’s slow oscillations often proceed relatively independently of frontal cortex, supporting the notion that anesthetics do not suppress the cortex uniformly but rather produce a spatial gradient in which posterior areas can still cycle in and out of transient Up states. Transfer entropy analyses further reinforce a feedforward-like (PPC→PFC) information flow under sevoflurane, contrasting with the robust bidirectional connectivity expected in wakefulness. These findings echo other reports in rodents and ferrets where the frontal cortex shows preferential inhibition while posterior regions retain partial or disjointed excitability (Raz et al., 2014; Sellers et al., 2013). Spatiotemporal pattern of the firing during Up states in PPC indicated that these transient Up states originated in deeper cortical channels, presumably layer V, consistent with layer V’s tendency to initiate bursts under sedation. Laminar recordings in rodents and cats demonstrate a clear deep-to-superficial progression at Up-state onset (Chauvette et al., 2010; Perez-Zabalza et al., 2013), and studies attribute this to the unique burst-generating ability and extensive connectivity of layer V pyramidal neurons (Lőrincz et al., 2015).

While these results deepen our understanding of sevoflurane-induced changes in frontoparietal networks, several limitations must be considered. The monkeys involved in this study were relatively old (21 and 18 years), which can affect both anesthetic metabolism and cortical reactivity. In addition, the probe locations were only putative and not verified histologically, making it difficult to definitively assign spiking activity to specific laminar compartments. Future investigations would benefit from postmortem tissue analysis or complementary imaging techniques to clarify laminar boundaries, along with broader spatial coverage (e.g., temporal or occipital recordings) to determine whether the feedforward-like phenomena observed here generalize across cortex. Exploring multiple anesthetic agents (such as propofol, dexmedetomidine, or ketamine) could reveal whether our findings on slow oscillations, gamma surges, and frontoparietal disconnection are agent-specific or reflect a broader principle of cortical inactivation. Finally, within-subject repeated sessions and richer physiological markers may help to disentangle individual variability from universal dose–response mechanisms, ultimately refining our understanding of how frontoparietal networks disintegrate under anesthesia.

## 5. Conclusion

In summary, our recordings revealed a global suppression of neural activity under increasing sevoflurane concentrations in both monkeys, with PFC displaying notably greater spiking suppression than PPC. This effect was more pronounced in the older animal (M2), for whom mid-range anesthetic levels (3% and 4%) clustered with deeper anesthesia (6%), whereas in the younger animal (M1) those same concentrations resembled lighter anesthesia (2%). Despite uniformly diminished activity across cortical layers, up states reliably began in deeper layers (likely layer V) but failed to propagate widely throughout the brain, indicating that residual activity remained largely localized. Overall, the findings suggest that anesthesia does not uniformly suppress different cortical areas and emphasize a dose-dependent but spatially consistent suppression of cortical dynamics, indicating how large-scale integration are necessary for sustained cortical processing. This implies that anesthesia does not just “switch off” the brain but rather reconfigures its functional architecture into a predictable but somewhat flexible low-frequency regime. The observed fronto-parietal gradient of suppression underscores the sensitivity of PFC to anesthetic-induced inactivation, shows importance of developing region specific metrics as potential markers for more precise anesthetic monitoring in clinical and research settings.

